# A Comparative Study of Protein Structure Prediction Tools for Challenging Targets: Snake Venom Toxins

**DOI:** 10.1101/2023.05.05.539526

**Authors:** Konstantinos Kalogeropoulos, Markus-Frederik Bohn, David E. Jenkins, Jann Ledergerber, Christoffer V. Sørensen, Nils Hofmann, Jack Wade, Thomas Fryer, Giang Thi Tuyet Nguyen, Ullrich auf dem Keller, Andreas H. Laustsen, Timothy P. Jenkins

## Abstract

Protein structure determination is a critical aspect of biological research, enabling us to understand protein function and potential applications. Recent advances in deep learning and artificial intelligence have led to the development of several protein structure prediction tools, such as AlphaFold2 and ColabFold. However, their performance has primarily been evaluated on well-characterised proteins, and comparisons using proteins with poor reference templates are lacking. In this study, we evaluated three modelling tools on their prediction of over 1000 snake venom toxin structures with no reference templates. Our findings show that AlphaFold2 (AF2) performed the best across all assessed parameters. We also observed that ColabFold (CF) only scored slightly worse than AF2, while being computationally less intensive. All tools struggled with regions of intrinsic disorder, such as loops and propeptide regions, and performed well in predicting the structure of functional domains. Overall, our study highlights the importance of exercising caution when working with proteins that have poor reference templates, are large, and contain flexible regions. Nonetheless, leveraging computational structure prediction tools can provide valuable insights into the modelling of protein interactions with different targets and reveal potential binding sites, active sites, and conformational changes, as well as into the design of potential molecular binders for reagent, diagnostic, or therapeutic purposes.

**Statement:** Recent advances in machine learning have led to the development of new protein structure prediction tools. However, these tools have mainly been tested on well-known proteins and their performance on proteins without known templates is unclear. This study evaluated the performance of three tools on over 1000 snake venom toxins. We found that while caution is required when studying poorly characterised proteins, these tools offer valuable opportunities to understand protein function and applications.

## Introduction

Understanding 3-dimensional (3D) protein structures is key to many research questions, ranging from fundamental topics concerning how a given protein functions to translational hurdles involving using such information to manipulate protein function for industrial, therapeutic or other purposes. The most reliable approach towards resolving protein structures has been the use of experimental technologies, such as x-ray crystallography, and more recently, cryogenic electron microscopy. Yet, such approaches are time-consuming, low-throughput, and sometimes impossible for difficult to crystallise targets such as membrane proteins (Kermani, 2021). In an attempt to increase throughput and allow for 3D characterisation of large structural datasets and produce structures of proteins where conventional approaches fail, scientists have explored computational methods for the prediction of structures based on evolutionary history. Homology modelling aims to predict the 3D structure of a given protein (target) sequence based on its homology to solved structures (templates) (Bai et al., 2015; Thompson et al., 2020) and pairwise evolutionary correlations (Altschuh et al., 1987; Jones et al., 2012; Marks et al., 2011; Shindyalov et al., 1994; Weigt et al., 2009). Homology modelling has been used to produce structure models for at least one domain in more than half of all known sequences and a total of over 38 million models deposited on ModBase (Pieper et al., 2014). The majority of these deposited structures were generated using the software program MODELLER (MDLR), an established and excellent protein modelling structure tool (Bitencourt-Ferreira & de Azevedo, 2019; Webb & Sali, 2016). This approach has aided in our understanding of protein structures and has grown in accuracy with the increasing number of experimental structures deposited in the Protein Data Bank (PDB) (“Protein Data Bank: The Single Global Archive for 3D Macromolecular Structure Data,” 2019), the rise of genomic sequencing, and the availability of deep learning techniques allowing rapid interpretation of these data. Nevertheless, contemporary evolutionary-history-based approaches, more often than not, fall short of generating predictions comparable to experimental accuracy. Predictions are particularly poor for proteins without close and resolved homologues. This has limited the utility of homology protein structure modelling for many biological applications to date.

Recently, protein structure prediction has, however, undergone a renaissance with the application of sophisticated machine learning approaches. Whilst these algorithms are still reliant on template proteins and multiple sequence alignment (MSA), the shift from a rational decision tree to transformer-based neural networks has seen a substantial improvement in prediction accuracy (Jumper et al., 2021). Indeed, the first protein structure prediction tool using transformers, i.e. AlphaFold2 (AF2) (Jumper et al., 2021), achieved the highest accuracy prediction at the Critical Assessment of protein Structure Prediction (CASP) competition (CASP14) (Pereira et al., 2021, p. 14). Its accuracy was comparable to experimental protein structure determination with 36% of their submitted protein targets having a root-mean-square deviation (RMSD) under 2 Å (generally considered to be a solved structure), and 86% under 5 Å, with a total mean of 3.8 Å which presented an impressive performance comparable to experimental accuracies. Since then, AF2 predicted structures for the near-whole proteome of 48 species with over 200 million entries (https://alphafold.ebi.ac.uk/) have been made publicly available, and the transformer-based approach of AF2 has also been independently reproduced in another tool,RoseTTAFold (Baek et al., 2021).

Whilst the prediction precision of these tools is unprecedented, the computational power required to leverage them is substantial. Primarily, building the MSAs is computationally intensive (multiple hours and >2TB of storage per protein) and involves sensitive homology searches via HMMER (Eddy, 2011) and HHblits (Steinegger et al., 2019). Further, running the deep neural networks for the modelling itself also requires computational power and memory, albeit negligible compared to building the MSAs. With this need for substantial computation in mind, researchers developed ColabFold (CF) (Mirdita et al., 2021), which harnesses Google Colaboratory and thus provides free access to powerful graphics processing units (GPUs). CF also accelerates predictions (20-30 times faster than AF2) by using MMseqs2 search (Mirdita et al., 2019; Steinegger & Söding, 2017) instead of AF2’s native input feature generation. Further, CF also leverages optimisation strategies for predictions of multiple structures by avoiding recompilation and adding early stop criteria. Still, the question remains of what approach to rely on, if any, for protein structure prediction. The reliance of MDLR, AF2, and CF on homology alignments suggests that whilst all of these approaches excel in well-characterised areas of biology, they might struggle when few high quality templates exist. This is the case for snake venom toxins. Here, 19,000-25,000 toxins are predicted to exist (Laustsen et al., 2016), but only around 2000 different proteins have been described, and an even smaller percentage (<10%) of their structures have been experimentally resolved. Yet, understanding venom toxin structures could carry many benefits in either harnessing their beneficial potential as therapeutics (Ferraz et al., 2019; Li et al., 2018; Mohamed Abd El-Aziz et al., 2019) or for developing better treatments for snakebite envenomings (Gutiérrez et al., 2017; Jenkins et al., 2019; Knudsen et al., 2018).

Thus, to investigate the performance of three commonly used protein structure prediction tools, we predicted the structures of 1062 snake venom toxins using MDLR and CF and compared them to each other and to AF2 predicted structures.

## Results

### Generation, retrieval, and validation of toxin structures

For this study, we retrieved 1062 snake venom toxin sequences from Uniprot, including 220 C-type lectins (CTLs), 82 disintegrins (DISs), 145 kunitz-type serine protease inhibitors (KUNs), 190 phospholipase A_2_s (PLA_2_s), 135 snake venom metalloproteinases (SVMPs), 147 snake venom serine proteases (SVSPs), and 274 three-finger toxins (3FTxs). Structures were generated for all of these sequences using MDLR, as well as CF. The respective AF2 structures were retrieved from the database. For any structures that had a propeptide, it was trimmed to allow for an equal comparison across tools. Notably, all tools appeared to model these propeptides very poorly. The 3186 trimmed structures were evaluated via a series of parameters, including Clash scores, MolProbity scores, Ramachandran favoured percentage and Ramachandran outlier percentage. AF2 was significantly better than both MDLR and CF across all scores. MDLR performed the worst, though it was only slightly worse than CF in both Ramachandran evaluations (Fig. 1; Table S1). Notably, MDLR also performed worse on Clash and MolProbity scores, the more amino acid residues a given structure had (Fig. S2).

**FIGURE 1.**
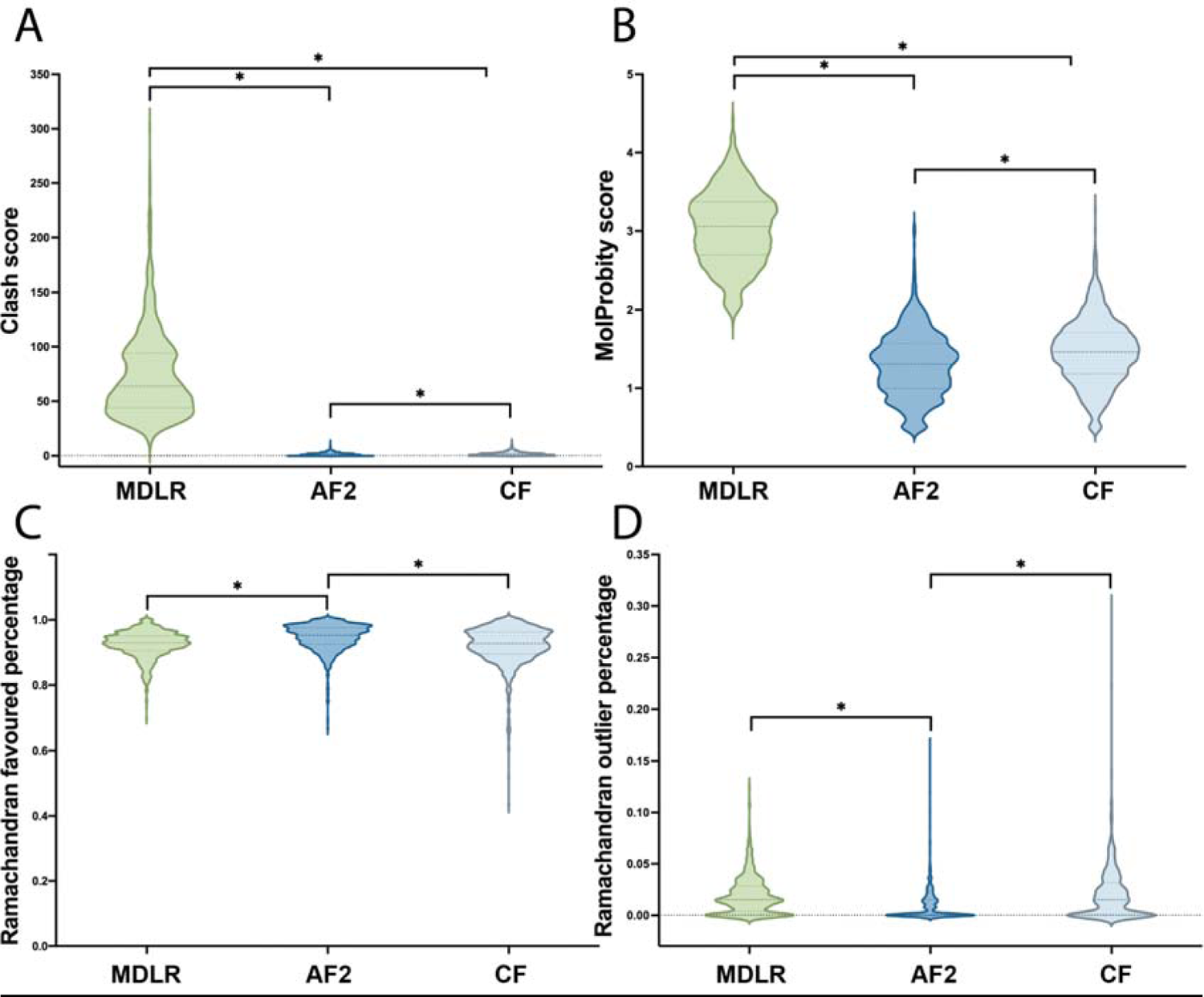
Quality evaluation of the toxin structures predicted by Modeller (MDLR), AlphaFold2 (AF2), and ColabFold (CF). A) Clash scores, *i.e.*, the number of serious clashes per 1000 atoms, defined as all non–donor–acceptor atoms overlapping by more than 0.4 Å. B) MolProbity scores (scores measured in percentiles; percentile ≥ 66 being the best); Ramachandran outliers (scores from 0 to 1; 1 being the best); Ramachandran favoured percentage (scores from 0 to 1; 1 being the best). Significant differences were established via Wilcoxon matched-pairs signed rank test and indicated by an asterisk (P<0.05).

### Comparing all Modeller, ColabFold, and AlphaFold 2 structures

To understand the differences in the structure prediction of the three modelling tools, all of the toxin structures underwent a three-way comparison. The largest variation in RMSD between models was observed in SVMPs, whereas the smallest was found to be for KUNs (Fig. 2; Table S3). RMSD was significantly different across all toxin families between AF2/MDLR and CF/MDLR, when compared to AF2/CF. Meanwhile, differences between AF2/MDLR and CF/MDLR were insignificant for all toxin families besides CTLs and 3FTxs. This indicated that AF2 and CF models were much more similar to each other than to MDLR generated structures. The largest differences between AF2 and CF models were found within the CTL and SVMP families.

**FIGURE 2.**
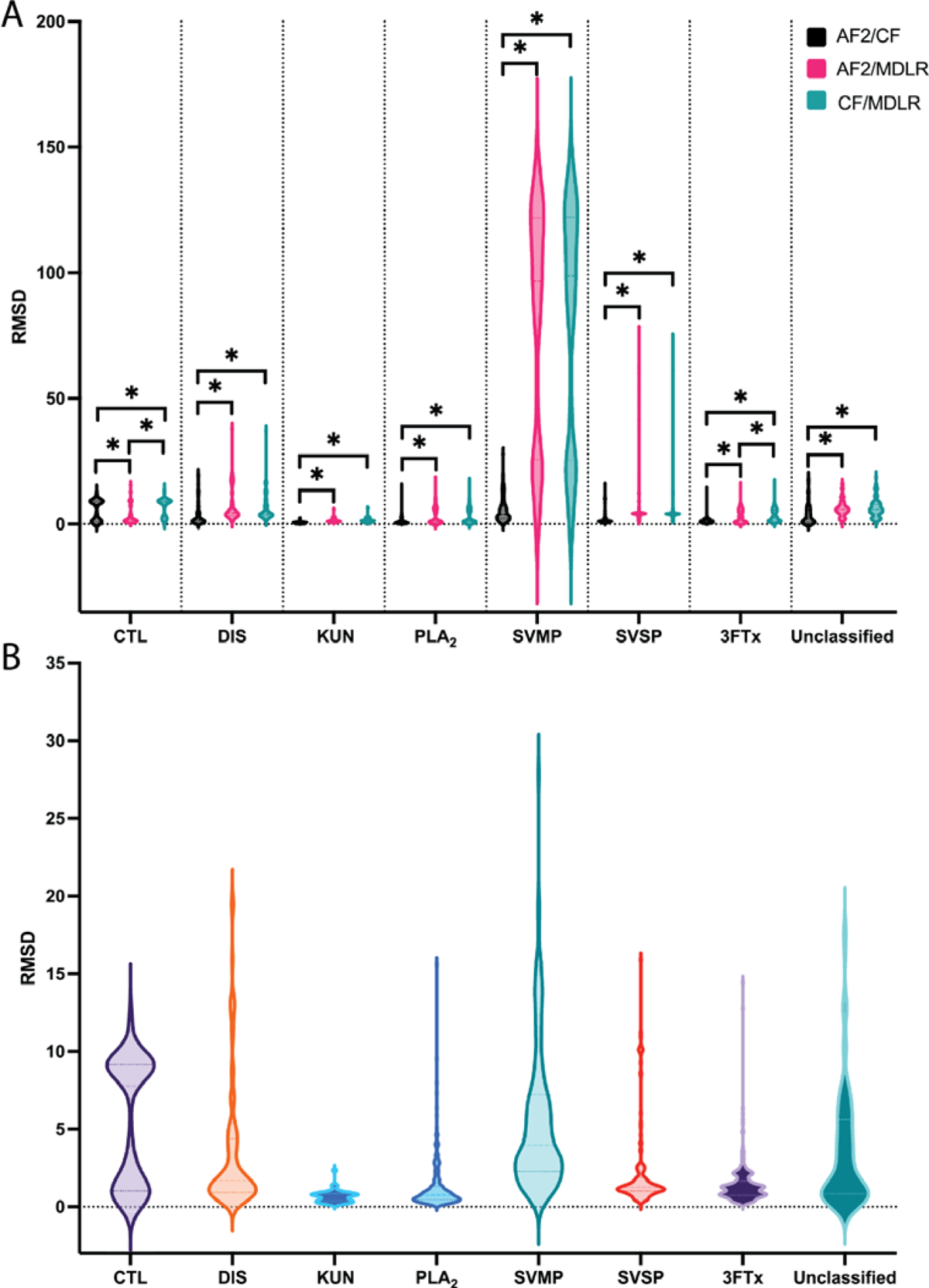
Differences in root-mean-square deviation (RMSD) between the toxin structures predicted by Modeller (MDLR), AlphaFold2 (AF2), and ColabFold (CF) and across the following toxin families: C-type lectins (CTLs), disintegrins (DISs), kunitz-type serine protease inhibitors (KUNs), phospholipase A_2_s (PLA_2_s), snake venom metalloproteinases (SVMPs), snake venom serine proteases (SVSPs), and three-finger toxins (3FTxs). A) This resulted in three comparisons, *i.e.,* AF2/CF, AF2/MDLR, and CF/MDLR. Significant differences were established via Wilcoxon matched-pairs signed rank test and indicated by an asterisk (P<0.05). B) Differences between AF2 and CF only.

Across all tools, it was found that the mean difference in RMSD was 14.80 Å with a standard deviation of 30.94 Å between MDLR and CF, 14.44 ± 30.97 Å between MDLR and AF2, and 2.77 ± 0.08 Å between AF2 and CF (Table S3). The highest similarity across all three was observed between models of toxins sharing the classic three-finger toxin fold, such as in P60774 from *Naja samarensis*, with an RMSD of 0.4 Å between MDLR and AF2 and no appreciable differences (Fig. 2A). On the other hand, the largest difference observed by RMSD (145 Å) was between the metalloproteinase Q10749 model generated by MDLR and either model generated by CF or AF2 (*Naja mossambica*). Modeller and AlphaFold agree only on the core of the peptidase domain, with AF2/CF identifying additional N- and C-terminal domains, which remain unstructured in the MDLR model (RMSD 145 Å between MDLR and both AF2 and CF; 3 Å between AF2 and CF). Restricting the alignment to the peptidase domain leads to a much better fit between models (RMSD 0.4/0.42 Å between MDLR and both AF2 and CF; 0.29 Å between AF2 and CF; Fig. 2B).

### Toxin family specific patterns

Snake venom toxins comprise a plethora of protein families substantially varying in size and structural complexity (Tasoulis & Isbister, 2017). Therefore, we also explored the predictions across all represented protein families.

### C-type lectin (CTLs)

Snake venom CTLs are 10-30 kDa glycoproteins that contain conserved carbohydrate recognition domains and can bind to specific sugar residues, resulting in various biological effects (Oliveira et al., 2022). Overall, we observed a mean RMSD difference of 5.2 ± 2.6 Å between all CTL models. The highest similarity was found to be between D8VNS6 from *Cerberus rynchops* (RMSD 0.4 Å between AF2 and CF, 0.9/08 Å between Modeller and AF2/CF) (Fig. 4A). The largest difference was between Q6X5S5 from *Echis ocellatus* (8.2 Å between AF2 and CF, as well as 15.7 Å and 13.6 Å differences between Modeller and AF2/CF, respectively; Fig. 4B). The largest difference between AF2 and CF predictions was found to be for A7X3W1 from *Pseudoferania polylepis* (12.6 Å; Fig. 4C). RMSD was significantly different between all three comparisons, with AF2/MDLR having the lowest average RMSD (3.3 Å) and CF/MDLR the highest (6.9 Å; Fig. 4D).

**FIGURE 3.**
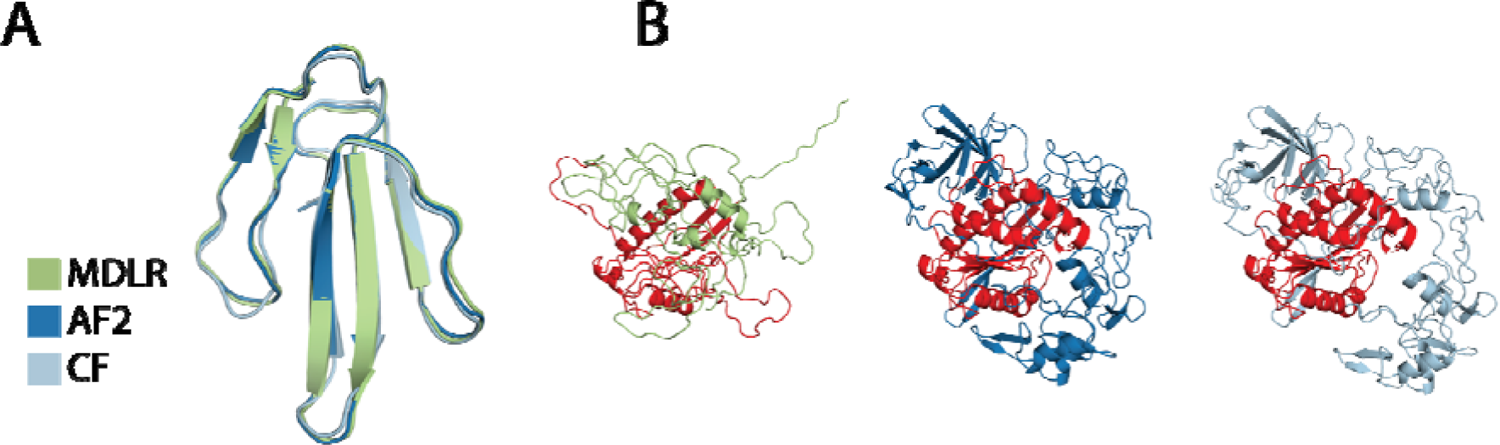
The structures predicted by Modeller (MDLR), AlphaFold2 (AF2), and ColabFold (CF), with the (A) most (P60774) and (B) least (Q10749) overlap across the entire dataset. Peptidase domain used for alignment used in (B) highlighted (red).

**FIGURE 4.**
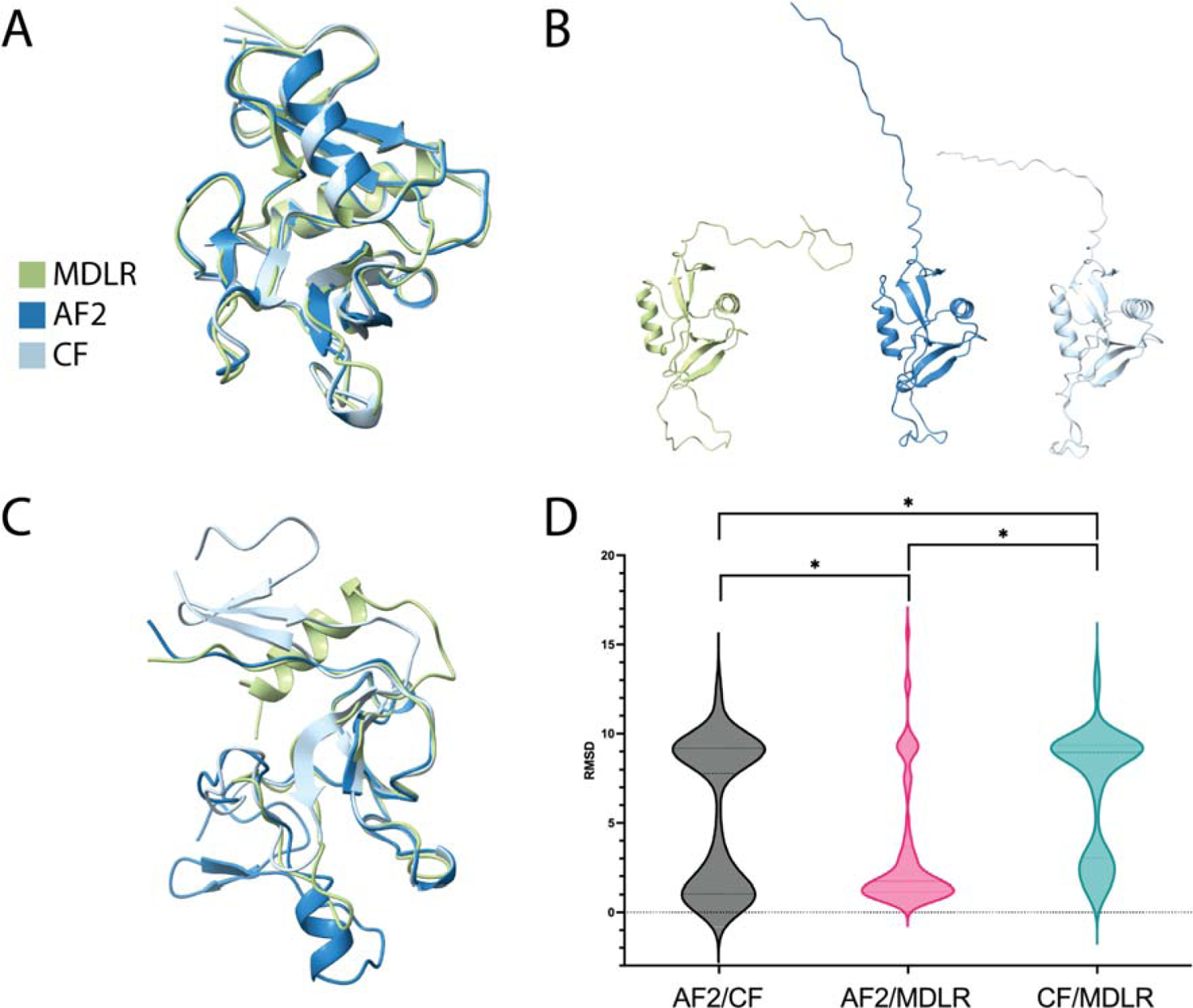
The structures predicted by Modeller (MDLR), AlphaFold2 (AF2), and ColabFold (CF), with the (A) most (D8VNS6) and (B) least (Q6X5S5) overlap across C-type lectins. (C) The largest difference between AF2 and CF predictions (A7X3W1). D) Differences in root-mean-square deviation (RMSD) between the toxin structures predicted by MDLR, AF2, and CF. Significant differences were established via Wilcoxon matched-pairs signed rank test and indicated by an asterisk (P<0.05).

### Disintegrins (DISs)

Snake venom disintegrins are non-enzymatic proteins that originated from SVMPs (Oliveira et al., 2022). Overall, we observed a mean RMSD difference of 7.0 ± 6.3 Å between MDLR and AF2 models. For example, the disintegrin fold of EC3B from *Echis carinatus* is predicted under good agreement (RMSD 4.9 Å between AF2 and CF, as well as 1.1/1.0 Å between MDLR and AF2/CF Fig 5A), with all solutions showing the same conserved four disulfide bonds expected from the heterodimeric disintegrin fold family. Here, most of the differences arise from the terminal peptidic regions that are not part of the disulfide-linked core. The largest differences were found between P0DJ43 models (RMSD 38/37 Å between MDLR and AF2/CF, 20 Å between AF2 and CF) from *Micropechis ikaheka* (Fig. 5B). There is very little agreement across the three different models. Notably, the only section scored by AlphaFold to have “very high confidence” (pLDDT > 90) is a < 20 amino acid residue segment within the disintegrin domain. All three tools were able to model the predicted disulfide bonding pattern. The largest difference between AF2 and CF predictions was found to be for P0DJ43 (19.7 Å; Fig. 5C). RMSD was significantly different between AF2/CF and both AF2/MDLR and CF/MDLR, with AF2/CF having the lowest average RMSD (3.6 Å) and AF2/MDLR the highest (7 Å; Fig. 5D).

**FIGURE 5.**
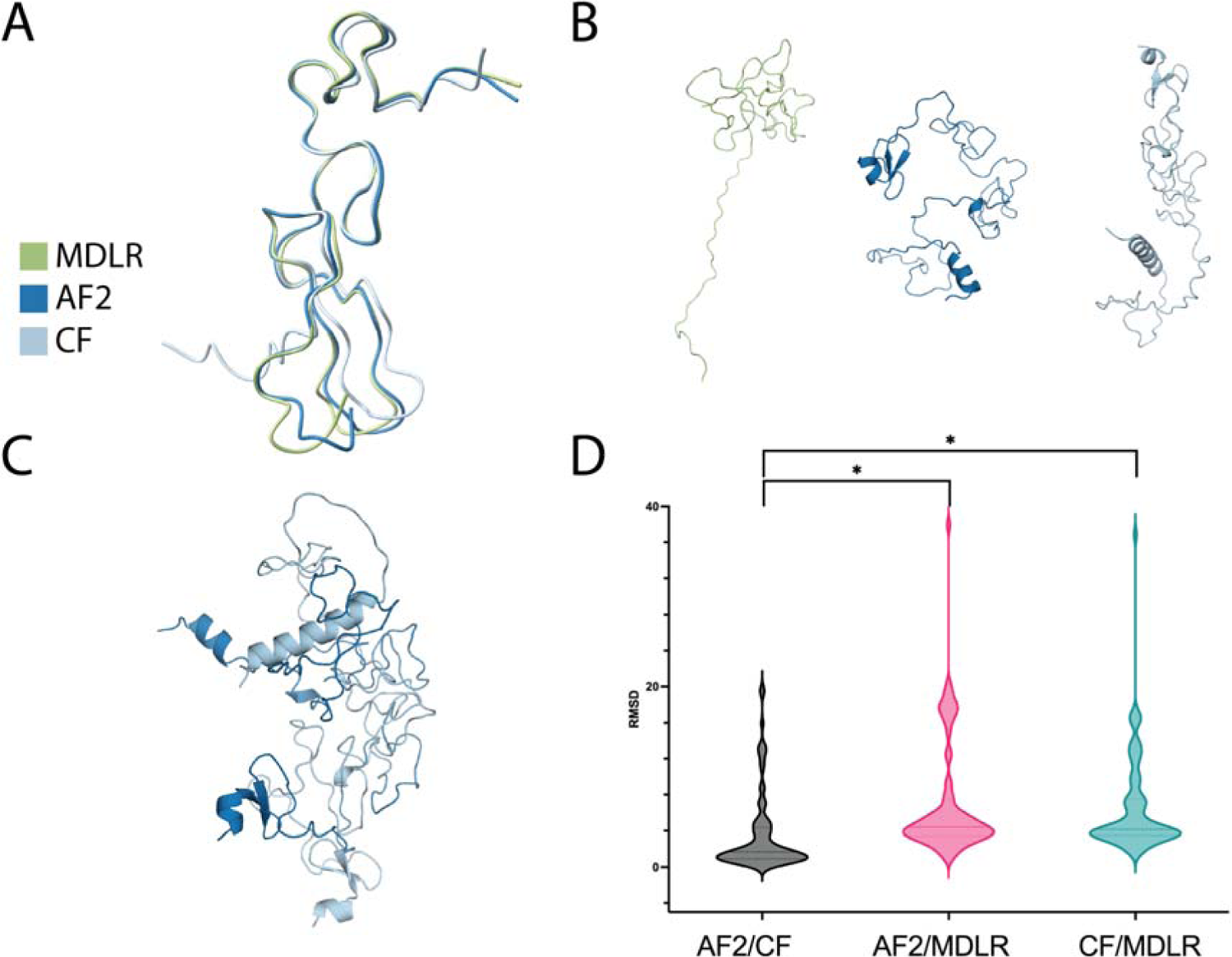
The structures predicted by Modeller (MDLR), AlphaFold2 (AF2), and ColabFold (CF), with the (A) most (P81631) and (B) least (P0DJ43) overlap across disintegrins. (C) The largest difference between AF2 and CF predictions (P0DJ43). D) Differences in root-mean-square deviation (RMSD) between the toxin structures predicted by MDLR, AF2, and CF. Significant differences were established via Wilcoxon matched-pairs signed rank test and indicated by an asterisk (P<0.05).

### Kunitz-type serine protease inhibitors (KUNs)

KUNs are 6-7 kDa small proteins containing three unique disulfide bonds that can inhibit the proteolytic activities of serine proteases (Oliveira et al., 2022). Overall, we observed a mean RMSD difference of 2.8 ± 3.3 Å between all KUN MDLR and AF2 models. The highest similarity is between C1IC51 from *Walterinnesia aegyptia* (RMSD 0.2 Å between AF2 and CF, 0.6 Å between Modeller and AF2/CF) (Fig. 4A). The largest difference driven by the terminal loop regions is between H6VC05 from *Daboia russelii* (2.0 Å between AF2 and CF, as well as 6.0 Å and 6.4 Å differences between Modeller and AF2/CF, respectively; (Fig. 4B). The largest difference between AF2 and CF predictions was found to be for P0CAR0 (2.5 Å; Fig. 4C). RMSD was significantly different between AF2/CF and both AF2/MDLR and CF/MDLR, with AF2/CF having the lowest average RMSD (0.7 Å) and AF2/MDLR as well as CF/MDLR shared highest (1.6 Å; Fig. 6D).

**FIGURE 6.**
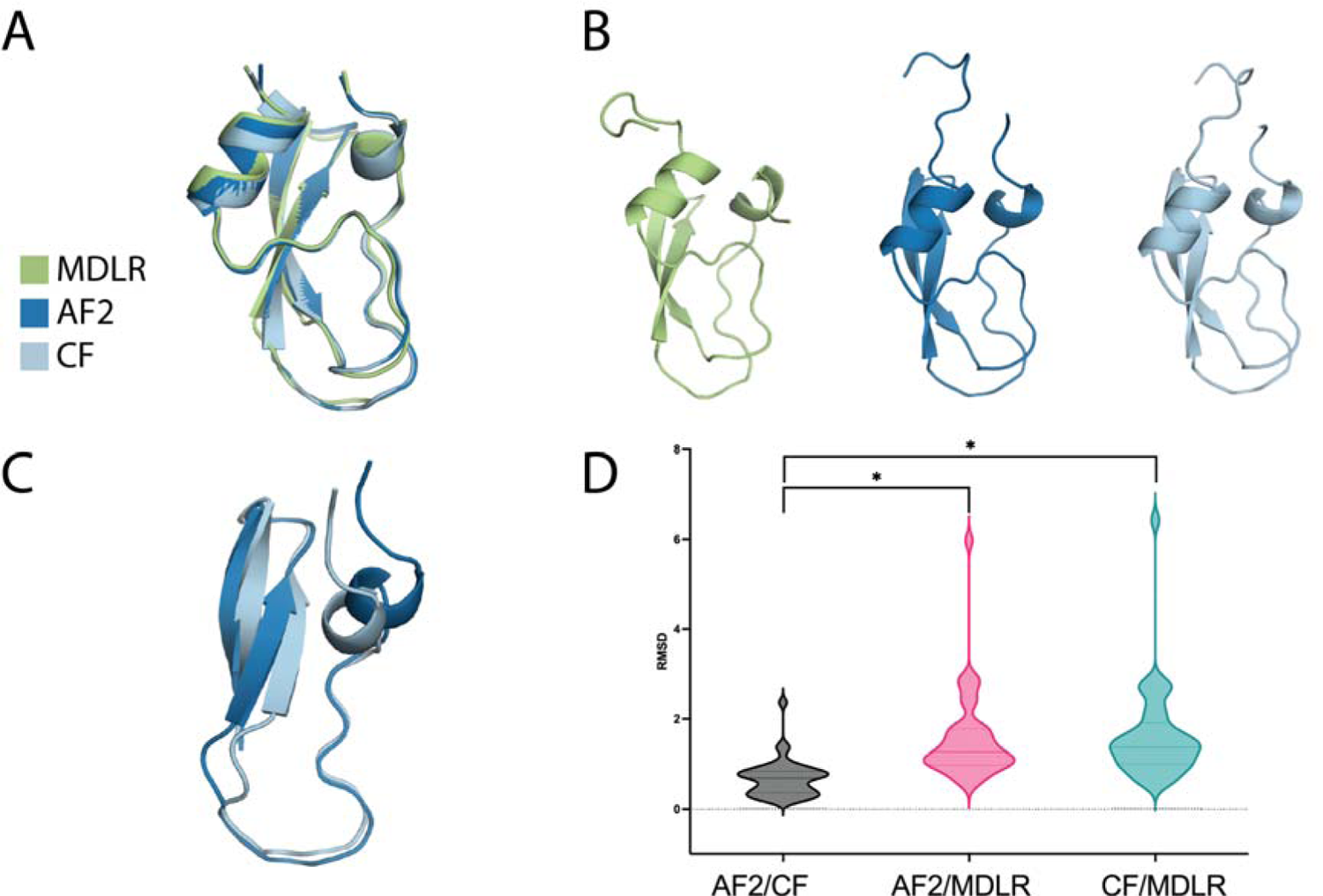
The structures predicted by Modeller (MDLR), AlphaFold2 (AF2), and ColabFold (CF), with the (A) most (C1IC51) and (B) least (H6VC05) overlap across kunitz-type serine protease inhibitors. (C) The largest difference between AF2 and CF predictions (P0CAR0). D) Differences in root-mean-square deviation (RMSD) between the toxin structures predicted by MDLR, AF2, and CF. Significant differences were established via Wilcoxon matched-pairs signed rank test and indicated by an asterisk (P<0.05).

### Phospholipase A_2_s (PLA_2_s)

Snake venom PLA_2_s are 13–19 kDa proteins and one of the main components of animal venoms (Oliveira et al., 2022). Overall, we observed a mean RMSD difference of 3.2 ± 3.4 Å between MDLR and AF2 models (Table 4). The highest similarity is detected for P04417 from *Gloydius blomhoffii* (with an RMSD of 0.9 Å between AF2 and CF, 0.4/0.9 Å between MDLR and AF2/CF) (Fig. 5A). The PLA_2_ domain itself is modelled to great agreement in all cases. The largest difference across predictions was found between P14411 from *Bungarus fasciatus* (5.2 Å between AF2 and CF, 5.8 Å between MDLR and AF2/CF), with differences mainly arising in positioning of the terminal loop regions (Fig. 5B). The largest difference between AF2 and CF predictions was found to be for Q8AY47 (15.6 Å; Fig. 5C). RMSD was significantly different between AF2/CF and both AF2/MDLR and CF/MDLR, with AF2/CF having the lowest average RMSD (1.4 Å) and CF/MDLR the highest (3.3 Å; Fig. 7D).

**FIGURE 7.**
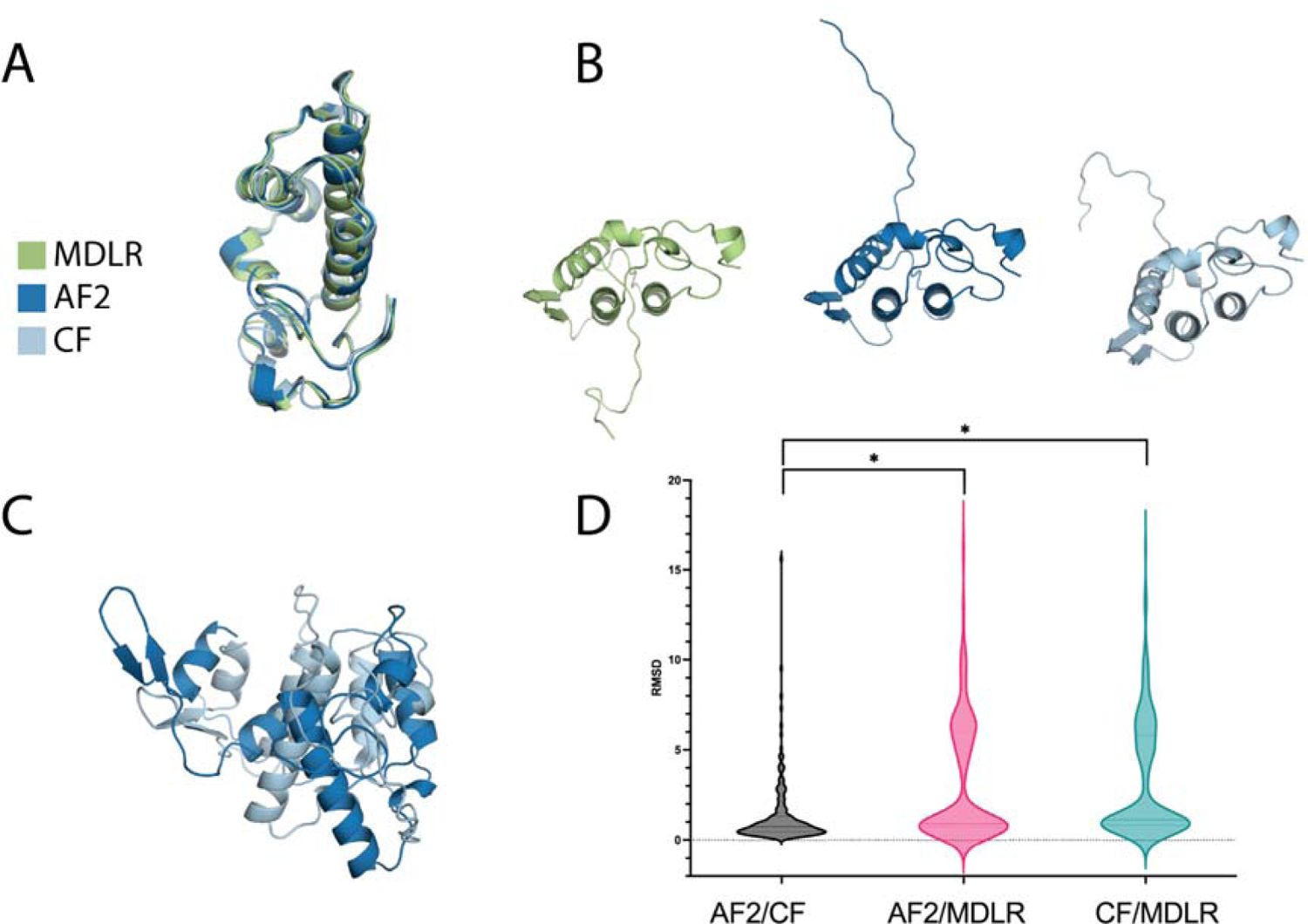
The structures predicted by Modeller (MDLR), AlphaFold2 (AF2), and ColabFold (CF), with the (A) most (P04417) and (B) least (P14411) overlap across phospholipase A_2_s. (C) The largest difference between AF2 and CF predictions (Q8AY47). D) Differences in root-mean-square deviation (RMSD) between the toxin structures predicted by MDLR, AF2, and CF. Significant differences were established via Wilcoxon matched-pairs signed rank test and indicated by an asterisk (P<0.05).

### Snake venom metalloproteinases (SVMPs)

SVMPs are a class of enzymes found in the venom of snakes, with sizes ranging from 20-100 kDa (Oliveira et al., 2022). Out of 137 SVMPs (average RMSD 80 ± 46 Å between MDLR and AF2 models), the highest similarity between models can be found in P20897 from *Crotalus ruber* (3.2 Å between AF2 and CF, 145 Å between Modeller and AF2/CF; Fig.6A). Meanwhile, the largest difference was observed between models of Q10749 from *Naja mossambica* (6.8 Å between AF2 and CF, 8.7 Å/ 7.8 Å between Modeller and AF2/CF; Fig. 6B) mentioned earlier. The largest difference between AF2 and CF predictions was found to be for Q3HTN1 (27.8 Å; Fig. 6C). RMSD was significantly different between AF2/CF and both AF2/MDLR and CF/MDLR, with AF2/CF having the lowest average RMSD (6.6 Å) and CF/MDLR the highest (81 Å; Fig. 8D).

**FIGURE 8.**
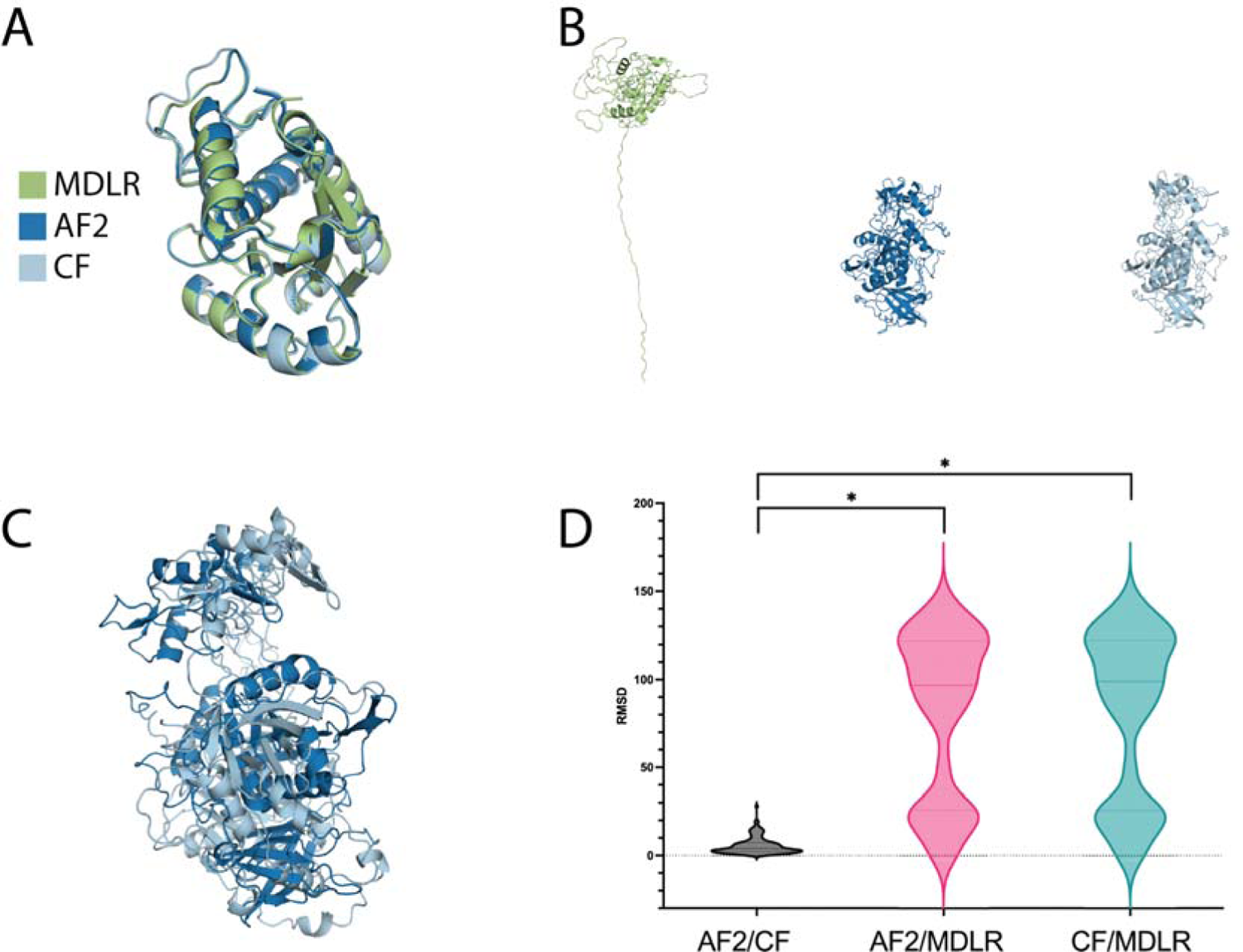
The structures predicted by Modeller (MDLR), AlphaFold2 (AF2), and ColabFold (CF), with the (A) most (P20897) and (B) least (Q10749) overlap across snake venom metalloproteinases. (C) The largest difference between AF2 and CF predictions (Q3HTN1). D) Differences in root-mean-square deviation (RMSD) between the toxin structures predicted by MDLR, AF2, and CF. Significant differences were established via Wilcoxon matched-pairs signed rank test and indicated by an asterisk (P<0.05).

### Snake venom serine proteases (SVSPs)

SVSPs are another class of enzymes found in snake venom, with sizes ranging from 26-67kDa (Oliveira et al., 2022). Out of the 147 serine proteases (average RMSD 8 ± 13 Å between MDLR and AF2 models), the largest similarity between models can be found for Q7SZE2, from *Gloydius ussuriensis* (0.7 Å between AF2 and CF, 0.5 Å and 0.9 Å between MDLR and AF2/CF respectively; Fig. 7A). The largest disagreement between tools can be found for Q58L94 from *Notechis scutatus* (10 Å between AF2 and CF, 78/75 Å between Modeller and AF2/CF respectively; Fig. 7B). Overall, the spread of RMSD between highest and lowest similarity models between MDLR and AF2 is 8.3 ± 14 Å. The biggest difference between AF2 and CF in RMSD for a given SVSP was found to be 16 Å (A6MFK8; Fig. 7C). RMSD was significantly different between AF2/CF and both AF2/MDLR and CF/MDLR, with AF2/CF having the lowest average RMSD (2.2 Å) and AF2/MDLR as well as CF/MDLR with the shared highest (8.3 Å; Fig. 9D).

**FIGURE 9.**
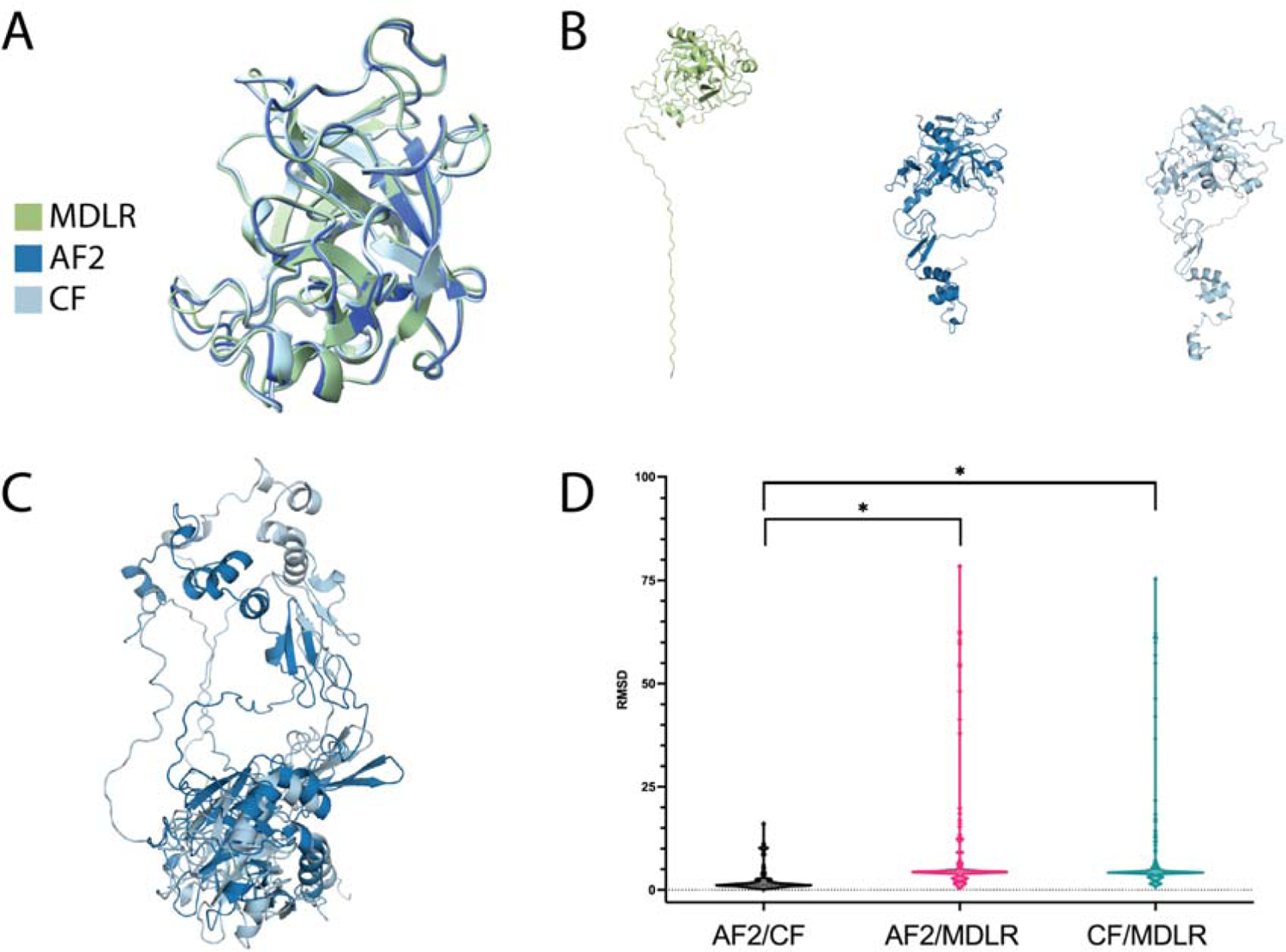
The snake venom serine protease structures predicted by Modeller (MDLR), AlphaFold2 (AF2), and ColabFold (CF), with the (A) most (Q7SZE2) and (B) least (Q58L94) overlap across snake venom serine proteases. (C) The largest difference between AF2 and CF predictions (A6MFK8). D) Differences in root-mean-square deviation (RMSD) between the toxin structures predicted by MDLR, AF2, and CF. Significant differences were established via Wilcoxon matched-pairs signed rank test and indicated by an asterisk (P<0.05).

### Three-finger toxins (3FTxs)

3FTxs comprise three major subfamilies of toxins, *i.e.,* cytotoxins, long-chain neurotoxins, and short-chain neurotoxins (Casewell et al., 2013). Out of the 275 3FTxs (average RMSD 3 ± 3 Å between MDLR and AF2 models), the greatest overlap between models was found to be for P60774 (0.37/0.92 Å between MDLR and AF2/CF, as well as 0.97 Å between AF2 and CF respectively; Fig. 8A). The largest difference in model RMSD was found to be for Q9W7K1 (12.8/12.7 Å between MDLR and AF2/CF, as well as 0.95 Å between AF2 and CF respectively; Fig. 8B). The protein found to have the largest differences in RMSD between AF2 and CF was P34074 (14.4 Å; Fig. 8C). RMSD was significantly different between all three comparisons, with AF2/MDLR having the lowest average RMSD (1.4 Å) and CF/MDLR the highest (3.5 Å; Fig. 10D).

**FIGURE 10.**
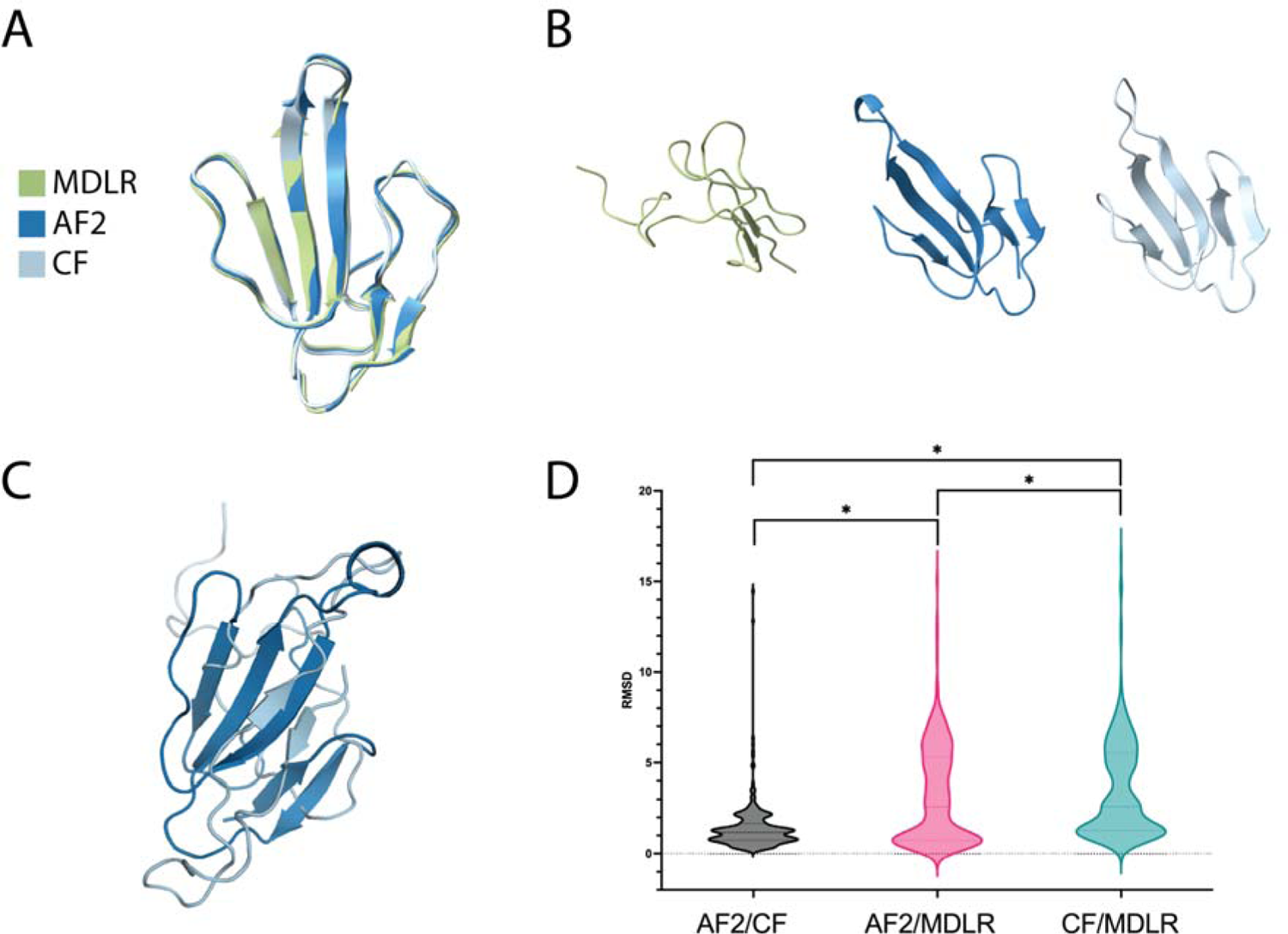
The three-finger toxin structures predicted by Modeller (MDLR), AlphaFold2 (AF2), and ColabFold (CF), with the (A) most (P60774) and (B) least (Q9W7K1) overlap. (C) The largest difference between AF2 and CF predictions (P34074). D) Differences in root-mean-square deviation (RMSD) between the toxin structures predicted by MDLR, AF2, and CF. Significant differences were established via Wilcoxon matched-pairs signed rank test and indicated by an asterisk (P<0.05).

## Discussion

Prediction of protein structures is a critical aspect of biological research, particularly in understanding the function and potential therapeutic applications of proteins or how protein-based disease targets can be targeted with therapeutic agents. With the advent of deep learning and AI, several protein structure prediction tools have emerged, such as Modeller (MLDR), ColabFold (CF), and AlphaFold 2 (AF2). Each of these tools utilise different algorithms and approaches, resulting in different predictions of protein structures. Therefore, in this study we aimed to provide insight into the reliability and accuracy of these three modelling tools using snake venom toxins as model proteins. A total of 1062 snake venom toxin sequences, representing seven protein families, were retrieved, and structures were generated for each sequence using MDLR and CF. The respective AF2 structures were retrieved from the database, and all 3186 structures (1062 from each tool) were evaluated using various parameters, including Clash score, MolProbity score, as well as Ramachandran favoured and outlier percentage to ensure no inherent quality bias was introduced by any of the tools. The results of the analysis revealed significant differences in performance across all three models, with the two non-homology-based approaches AF2 and CF exhibiting superior performance in all four evaluated parameters compared to the template-based method Modeller. As an additional observation, AF2 performed better than ColabFold across all evaluated parameters and significantly so in both Clash and MolProbity scores, which is in line with prior findings (Mirdita et al., 2021). These results indicate that non-homology-based approaches perform better on a range of different protein families found in snake venoms. Given that Modeller utilises a template-based approach, it is conceivable that its poor performance may be attributed to the lack of reported snake toxin structures.

To have a better understanding of the differences in structure prediction quality of the three modelling tools, a three-way comparison was conducted over all assessed 1062 snake toxins. RMSD analysis clearly indicated that a substantial spread existed in model overlap across the different toxins, ranging from 0.14–145 Å. The differences between tools were again greatest when MDLR was involved, whereas differences between AF2 and CF were rarely significant. Notably, when considering different snake toxin families, MDLR exhibited the smallest performance differences in short and highly conserved proteins, such as 3FTxs that share the classic three-finger toxin fold. MDLR is typically not suitable for domains that do not belong to a fold family because it is designed to work with targets where it can successfully assign a fold (Webb & Sali, 2016), and differences may arise in the twilight zone of short sequences with relatively low similarity (Khor et al., 2015). Consequently, the largest performance differences were observed for long and complex proteins, such as SVMPs. Nevertheless, MDLR was able to successfully model conserved functional cores in SVMPs, such as the peptidase domain. This is likely due to the high level of structural conservation of such sites due to the need to also conserve function (Alberts et al., 2002), but these observations also suggest that the performance of MDLR is heavily reliant on the quality and size of its template database. Another area where disagreement was observed between the generated protein structures was in loop regions. Whilst loop regions were typically identified across all three models we tested, their conformations often differed across MDLR, AF2, and CF. This was unsurprising and stems from proteins being dynamic molecules with a large conformational plasticity, allowing them to perform complex biological functions (Mukhopadhyay, 2022). Yet, these features are not uniformly distributed across the molecule, but are usually localised to parts with larger degrees of kinematic freedom (Papaleo et al., 2016). Thus, modelling conformations of loop regions remains challenging in computational biology and is usually inversely related to loop length (Barozet et al., 2021); these dynamics of flexibility, but also lack of reference structures, explains why we observed poor modelling of propeptides. It also highlights the need for a better structural understanding, due to propeptides’ key roles in chaperoning and to provide insights into how the interactions of the propeptide region inhibits enzymatic function for the design of inhibitor molecules. Finally, it is notable that the overall mean difference between AF2 and CF was close to 3 Å, which is large enough to have a substantial impact when used for generative ML approaches for protein design (Wang et al., 2021; Watson et al., 2022). For some SVMPs, these differences even exceed 27 Å, highlighting the current limitations of the explored protein structure prediction tools (Bryant et al., 2022).

Nevertheless, several of the toxin families assessed in this article have structural models that are of sufficiently high quality for further analysis, and thus a multitude of use cases; specifically the highly conserved 3FTxs, KUNs, and PLA_2_s can be used for computational simulations to model their interactions with different targets. This can help reveal the details of their binding sites, active sites, and conformational changes, as well as for the discovery and design of potential molecular binders for reagent, diagnostic, or therapeutic purposes (Norman et al., 2020). Notably, even toxin families with higher model variability and uncertainty (i.e. DISs, SVSPs, and SVMPs) could be used for similar purposes, as long as the focus falls on their conserved functional domains, such as their active sites. Importantly, these considerations regarding the limitations of computational structural modelling are not only relevant for toxin researchers, but can also be transferred to any other research area that is using computational predictions for protein structures and particularly ones with poor experimental coverage, such as rare diseases (Rossi Sebastiano et al., 2022). Overall, we hope the unravelled dynamics of computational structure prediction and the provision of all models and their comparisons constitute a key resource that can help de-risk future analyses.

## Conclusion

The availability and accessibility of a range of powerful computational models are a game changer for structural biology. Whilst extremely powerful, the plethora of available tools each come with their own set of advantages and disadvantages; these are often somewhat understood within their application in model organisms, but little data exists on their performance on poorly characterised protein targets. Here, we studied the performance of three different computational structural biology tools on their predictions of over 2000 snake toxin structures that have few experimental reference structures available. We generated these structures using Modeller and ColabFold and compared them to each other as well as AlphaFold2 designs. We found that predictions were often closely aligned between CF and AF2, whereas MDLR often offered differing predictions. Nevertheless, differences between AF and CF were common, highlighting the need for cross-model validation of predicted structures. Notably, all tools performed well in predicting functional domains, while struggling with elements that are intrinsically disordered, such as loop regions. We further identified toxin families and structural features within these, as well as specific toxins, that were associated with substantially differing predictions across models. We therefore conclude that it is important to consider the complexity of the modelling task and use orthogonal modelling methods, such as AF2 and CF in combination with each other, to improve the reliability of structural assumptions. This will not only help future research quickly identify potential discrepancies and de-risk their use of these models, but also highlights key protein families, such as SVMPs, SVSPs, and DISs that require further experimental validation.

## Materials and methods

### Retrieval of snake venom toxin sequences

To retrieve the sequences of all published snake venom toxins (Nov 2020) belonging to potentially medically relevant toxin families, we used the utilities offered by VenomZone (*VenomZone*, n.d.). We selected C-type lectins (CTLs), disintegrins (DISs), kunitz-type serine protease inhibitors (KUNs), phospholipase A_2_s (PLA_2_s), snake venom metalloproteinases (SVMPs), snake venom serine proteases (SVSPs), and three-finger toxins (3FTxs), and searched for entries on uniprot (e.g., taxonomy:serpentes family:“phospholipase a2 family” (annotation:(type:“tissue specificity” venom) OR locations:(location:nematocyst)) AND reviewed:yes). The toxin Uniprot entry information was retrieved with custom Python scripts using Biopython’s (Cock et al., 2009) ExPASy package (Gasteiger et al., 2003).

### Identification of experimentally resolved toxin structures

The protein database (PDB) information on those entries was examined by cross-reference documentation in the obtained Uniprot files. Where there were multiple annotated PDB entries for a Uniprot accession, the entry with the highest resolution and chains corresponding to the toxin was used. This was achieved simply by parsing Uniprot files using custom scripts. Preference was given to X-ray structures. Thereafter, for a given toxin, the selected PDB file containing structural information for that entry was retrieved using PDB-tools (Rodrigues et al., 2018). Using the same tool, files obtained were filtered, selecting for chains corresponding to the toxin, and removing ligands and hydrogens in the structure. This process was also automated via bash scripting.

### Generating Modeller structures

For the remaining toxins that did not contain an annotated structure, the theoretical structure was predicted using homology modelling algorithms. To that purpose, we employed the Python MODELLER software (Webb & Sali, 2016). Toxins below the threshold of 30 amino acid length were discarded. Thereafter, the “multiple template modelling” approach and “loop refinement” was used with the PDB_95 database for template search. These methods significantly improve modelling quality, especially of complex regions, such as loops (Webb & Sali, 2016). Predictions were automated and run using multiple threading with subprocesses. For each toxin sequence, a total of 10 models were generated, and the best model, based on the lowest discrete optimised protein energy (DOPE) score, was selected as the representative model. The DOPE score (Shen & Sali, 2006) is based on an improved reference state that corresponds to non-interacting atoms in a homogeneous sphere with the radius dependent on a sample native structure; it thus accounts for the finite and spherical shape of the native structures. It is used to assess the energy of the protein model generated through many iterations by MODELLER. Finally, we evaluated the DOPE score of all of our toxin models to ensure the overall quality was sufficient to merit their usage for further research.

### Generating ColabFold structures

A version of ColabFold / MMSeqs2 (Steinegger et al., 2019) was modified to allow batch generation of structures. The program, originally designed to process single proteins via an online interface, was restructured in order to process multiple sequences from file input. Functions were rewritten to eliminate use of global variables, in order to permit loop processing. A mechanism was added to allow processing to be interrupted and restarted without repeating previously generated structures. This was essential to allow use of cheap compute facilities such as Google Colab and Colab Pro, which reserve the right to interrupt long-running jobs. This mechanism also allowed processing to be shared across multiple compute instances. The feature dictionary and all other parameters for reproducing the structure generation of any particular structure (and generation errors, if any) were written to sequence specific files during batch processing. The number of CF models calculated for each residue was set to the maximum, five; multi-sequence alignment was set; environmental processing was set; Amber relaxation was enabled; use of templates was enabled; and homooligomer processing was disabled. Query sequences of less than 20 residues or more than 1400 residues were skipped. Amber relaxation does not work for incomplete sequences with undetermined residues, so, Changes to the program were made to recognise these cases automatically and to invoke ColabFold without relaxation. The FASTA file processing was based on code from brentp (github) and Minkyung Baek’s modification to allow Colabfold to process complexes, as used in MMSeqs2, was retained (Baek & Baker, 2022).

### Retrieving AlphaFold2 structures

To retrieve relevant toxin structures predicted by AF2, we downloaded the latest database (https://alphafold.ebi.ac.uk/download) and selected the same list of UniProtIDs as were used in our MDLR and CF predictions.

### Trimming of propeptides

To enhance the accuracy and efficiency of our protein structure prediction we developed a code to detect and trim the signal peptides since these are not part of the mature protein or its function. The code is available via GitHub (https://github.com/nilshof01/signal_cleaver). Before starting the main script it is necessary to download the SignalP6 package. For the processing of the UniProtIDs and predicting potential signal peptides, we developed a Python 3.9 script to retrieve the sequences by giving a list of IDs as input. Next, the protein sequences are analysed using SignalP6, a state-of-the-art software capable of predicting the presence and location of all five signal peptide cleavage sites in protein sequences. The software generates a GFF (General Feature Format) file for each ID in a separate folder, which contains the start-, end position and score of the predicted signal peptide regions (Teufel et al., 2022). A further python script was developed to parse over the GFF files and extract this information for each ID in a pandas dataframe which was used to trim the sequences subsequently. Finally, the actual N-terminal trimming of the PDB file was performed using phenix.pdbtools (Liebschner et al., 2019). All structures have been deposited in Mendeley data (DOI: 10.17632/gjk47cjm26.1).

### Quality control and comparisons of structures

To assess the quality and validate the toxin structures predicted by MDLR, AF2, and CF, we used several different approaches. We evaluated the Ramachandran scores (outliers and favoured percentage), which indicate the number of amino acid residues with poor or favoured φ/ψ (Phi/Psi) angles. Phi/Psi angles are the dihedral angles in the protein backbone. Only certain angles are typically found in proteins. Finally, we compared our models’ Molprobity scores (Davis et al., 2007). MolProbity is a structure-validation web service which uses a weighted function of clashes, Ramachandran favoured, and rotamer outliers, scaled and normalised so that its value approximates the resolution at which that score would be average. To assess global and local differences between models generated by each of the three tools, we performed pairwise structural alignment of each toxin model using the superimpose method (Shindyalov & Bourne, 1998) from the Bio.PDB module in the Biopython toolbox and ranked aligned pairs by RMSD, separated by toxin family. Cases where RMSD was highest/lowest for each toxin family were inspected manually to identify areas of structural disagreement. Statistical analysis was performed as outlined above. To establish statistical significance the dataset was analysed via Prism (v.9.5.1). Once a lack of Gaussian distribution was established via both the Shapiro-Wilk and Kolmogorov-Smirnov test, the Wilcoxon matched-pairs signed rank test was selected as appropriate non-parametric analysis to conduct pairwise comparisons of AF2, CF, and MDLR.

## Supporting information

Table S1. Differences in root-mean-square deviation (RMSD) between the toxin structures predicted by Modeller (MDLR), AlphaFold2 (AF2), and ColabFold

Table S3. Differences in root-mean-square deviation (RMSD) between the toxin structures predicted by Modeller (MDLR), AlphaFold2 (AF2), and ColabFold

Fig. S2. Quality evaluation of the toxin models predicted by Modeller (MDLR), AlphaFold2 (AF2), and ColabFold (CF) by number of residues. A) Clash sco

## Supplementary files

**Table S1.** Differences in root-mean-square deviation (RMSD) between the toxin structures predicted by Modeller (MDLR), AlphaFold2 (AF2), and ColabFold (CF).

**Fig. S2.** Quality evaluation of the toxin models predicted by Modeller (MDLR), AlphaFold2 (AF2), and ColabFold (CF) by number of residues. A) Clash scores, i.e. the number of serious clashes per 1000 atoms, defined as all non–donor–acceptor atoms overlapping by more than 0.4 Å. B) MolProbity scores (scores measured in percentiles; percentile ≥ 66 being the best); Ramachandran outliers (scores from 0 to 1; 1 being the best); Ramachandran favoured percentage (scores from 0 to 1; 1 being the best).

**Table S3.** Differences in root-mean-square deviation (RMSD) between the toxin structures predicted by Modeller (MDLR), AlphaFold2 (AF2), and ColabFold (CF).

## Notes

### Competing Interest Statement

The authors have declared no competing interest.

