## Supplementary figures and images for "A Comparative Study of Protein Structure Prediction Tools for Challenging Targets: Snake Venom Toxins"

### Fig. S2. Quality evaluation of the toxin models predicted by Modeller (MDLR), AlphaFold2 (AF2), and ColabFold (CF) by number of residues. A) Clash sco

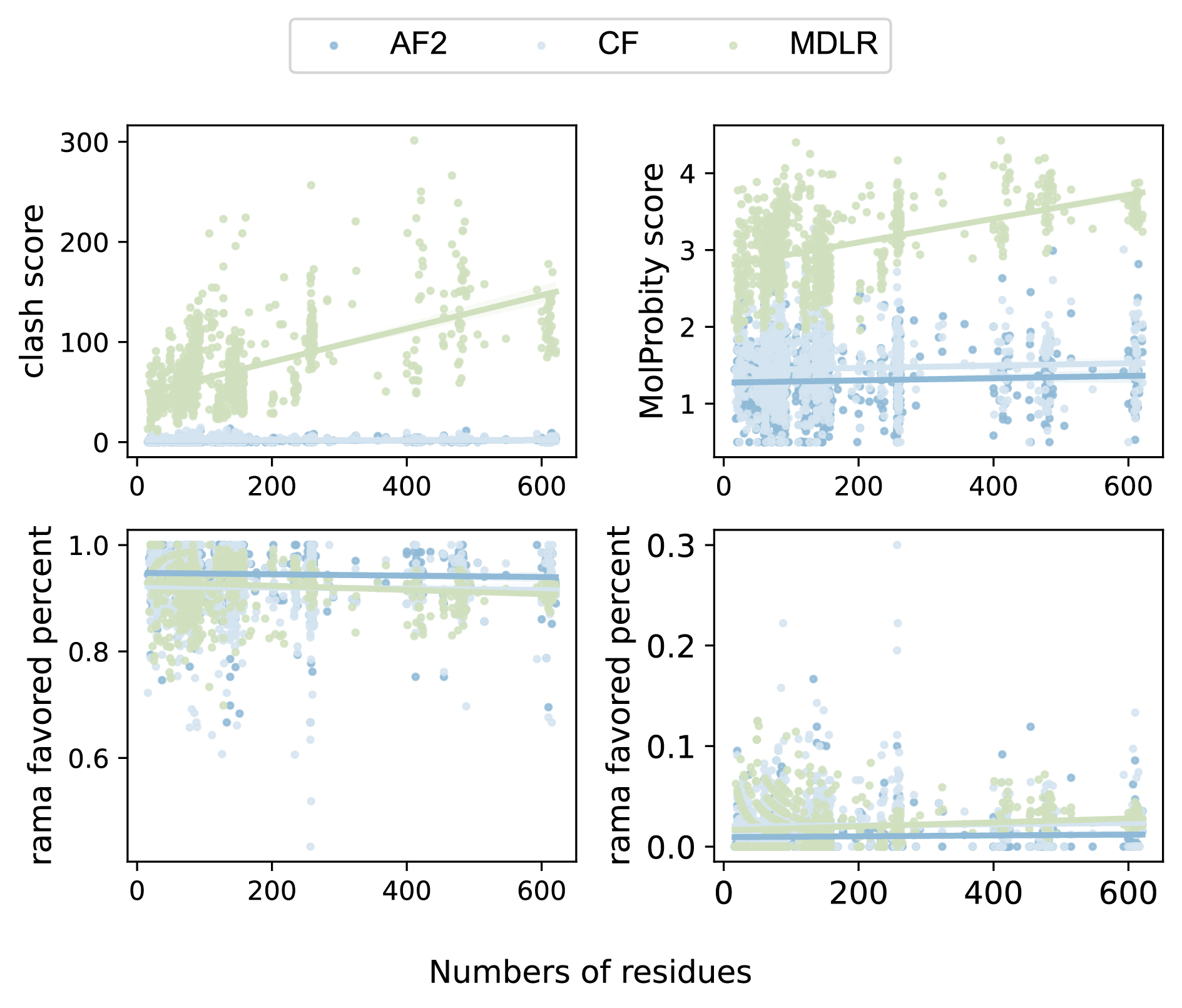
